# Stimulus Whitening Improves the Efficiency of Reverse Correlation

**DOI:** 10.1101/2022.01.25.477706

**Authors:** Alexis Compton, Benjamin W. Roop, Benjamin Parrell, Adam C. Lammert

## Abstract

Human perception depends upon internal representations of the environment that help to organize the raw information available from the senses by acting as reference patterns. Internal representations are widely characterized using reverse correlation, a method capable of producing unconstrained estimates of the representation itself, all on the basis of simple responses to random stimuli. Despite its advantages, reverse correlation is often infeasible to apply because the number of stimulus-response trials needed to provide an accurate estimate is typically very large. Prior approaches have aimed to overcome this sampling inefficiency by incorporating prior knowledge of the representation, which biases the estimate and ultimately limits the essential power of reverse correlation. The present approach, however, improves efficiency via stimulus whitening, a statistical procedure that decorrelates stimuli, making them less redundant, and commensurately more favorable for efficient estimation of an arbitrary target. We provide a mathematical justification for whitening, and demonstrate in simulation that whitening provides greater than 85% improvement in efficiency for a given estimation accuracy, and also a two- to five-fold increase in accuracy for a given sample size. Improving the efficiency of reverse correlation may enable a broader scope of investigations into individual variability and potential universality of perceptual mechanisms.

---

Reverse correlation is a powerful method for characterizing the underlying mechanisms of perception (De Boer & Kuyper, 1968; Ahumada & Lovell, 1971). It has a long history of use in characterizing the latent representations encapsulated in neural tuning (e.g., receptive fields; Ringach, 2004; Nishimoto, 2006), and has more recently become a primary method for inferring cognitive representations that drive the top-down processes of perception (e.g., face or phoneme recognition; Ahumada & Lovell, 1971; Gosselin & Schyns, 2001; Jäkel et al., 2009; Neri & Levi, 2006; Smith et al., 2012; Varnet et al., 2013), and even to estimate representations associated with abstract psychosocial categories (e.g., “male” vs. “female” faces; Brinkman et al., 2017; Mangini & Biederman, 2004; Moon et al., 2020; Ponsot et al., 2018). Indeed, the method has broad applicability for characterizing many aspects of neurological, cognitive, or psychological function (Ringach, 2004) and is closely related to the widely-used “white noise approach” to characterizing physiological (Marmarelis & Marmarelis, 1978) and engineering systems (Ljung, 1999).

In reverse correlation, stimulus-response data are elicited via the presentation of richly varying stimuli. For example, in psychophysical applications of reverse correlation, subjects may be presented with images composed of white noise and asked to make subjective “yes/no” responses about whether they perceived the presence of a specific signal, such as a face (e.g., Smith, 2012). Latent perceptual representations that optimally explain the pattern of responses can then be estimated by regressing subject responses against the stimuli over many trials, with the regression coefficients constituting an estimate of the representation itself.

However, current formulations of reverse correlation are widely known to be inefficient in the sense that many stimulus-response trials are required to achieve desirable estimation accuracies (Mineault, 2009). This inefficiency severely limits the feasibility of conducting reverse correlation studies to experimental protocols where subject participation can be maintained over extended timelines. Long protocols may mean that very few participants can be examined in any given study, and thus any analyses and inferences regarding possible universal aspects of human cognitive representation are severely limited. For example, in a notable study on representations of orthographic characters, Gosselin & Schyns (2003) collected 20,000 trials from three subjects over a period of two weeks. At the same time, inefficiency is an important consideration even for applications where collecting a large number of trials is feasible, because its existence implies that higher accuracies may be possible for a given number of trials if efficiency can be improved.

Attempts to improve efficiency of reverse correlation can be broadly characterized as either retrospective or prospective. *Retrospective* approaches impose some constraints on the inferred representations at the time of estimation, after data collection is complete. One common example of this approach is smoothing (e.g., low-pass filtering) the raw estimates (Gosselin & Schyns, 2003), which stems from the assumption that high-frequency information in the estimate is irrelevant noise. It has also been shown that the assumption of sparsity – i.e., that the target representation can be sparsely represented in some basis – can lead to dramatic improvements in efficiency when methods that incorporate this assumption are employed in the estimation process. For example, Mineault (2009) showed efficiency improvements using Generalized Linear Models with sparsity priors, and Roop (2021) employed a Compressive Sensing framework with L1 optimization.

On the other hand, *prospective* approaches to improving efficiency attempt to condition the stimuli in some way, prior to their presentation as part of the data collection. The most common example of this approach is to assume that the target has a certain form, even a very generalized one, and then construct stimuli that vary in relation to that form in specified ways. For example, approaches to psychosocial aspects of human faces (Mangini, 2003; Dotsch, 2012; Brinkman, 2017; Moon, 2020) have often proceeded from the assumption that representation of a trustworthy face is similar to a neutral face, and consequently generated stimuli by adding noise to an exemplar image of a neutral face. A similar approach has been taken in several auditory studies, in which stimuli were generated by adding noise to recordings of natural speech (Varnet 2013a; Varnet 2013b; Varnet 2015; Varnet 2016). Incorporating prior knowledge about the target representation into the stimuli improves efficiency by limiting variation along dimensions that are assumed to be irrelevant to the representation.

Whether retrospective or prospective, existing approaches to improving the efficiency of reverse correlation all function on the basis of some assumed knowledge regarding the target representation – i.e., that it has some general form, or is smooth or sparse – which is then incorporated either in the stimuli, in prospective approaches, or into the estimation process, in retrospective approaches. The assumed knowledge incorporated into existing approaches, even if well-justified, will exert a direct influence on estimates of the representation, limiting the essential power and promise of reverse correlation, which may be viewed as stemming from its ability to provide estimates that are unconstrained and unbiased. If, on the other hand, such assumed knowledge is not well-justified, then it will compromise the quality or interpretation of the estimate by introducing bias *a priori*.

Rather than relying on assumed knowledge regarding the target representation, the present work attempts, for the first time, to develop a prospective approach that instead conditions reverse correlation stimuli such that their general statistical properties are more favorable for efficient estimation of any arbitrary representation, without any *a priori* assumptions about the nature of that target. The present approach begins only with the knowledge that a randomly-generated set of stimuli will be expected to contain pairs of stimuli that are correlated by chance, especially under the conditions in which reverse correlation is typically applied – i.e., many stimuli that are low- to moderate-dimensional in size. Such correlation may be expected to decrease the effective sample size (Kish, 1969; Liang, 1993) of reverse correlation experiments using those stimuli by making observations overlapping and mutually predictable. Here, we prospectively employ *whitening*, a well-known statistical procedure for eliminating covariance in data, to improve the statistical properties of reverse correlation stimuli in this regard. We develop and present a mathematical justification for the effectiveness of stimulus whitening, and we demonstrate empirically that whitening can dramatically improve the efficiency of reverse correlation.

## Background

### Reverse Correlation

Reverse correlation follows, in essence, a regression model (see Fig 1). Subject responses, *y* ∈ {−1,1} are assumed to be generated by a process following

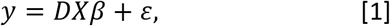

where the subject’ s internal representation is a p-by-1 vector *β*, the *n* stimuli are contained in a *n*-by-*p* matrix *X*, and *ε* is some noise. The matrix *D*, where *D_ii_* = 1/|*X_i_β*| for row *i* of *X*, acts to binarize the responses to values −1 (a negative response) and 1 (a positive response), equivalent to applying the signum function. Estimates of *β* can be obtained using the Normal equation,

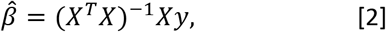

but are often made using the simplified formula

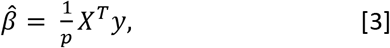

under the assumption that, across many stimuli, the columns of *X* will be nearly uncorrelated, meaning *X^T^X* = *I*, the identity matrix. Note that this assumption may become inappropriate for small values of *n*, for reasons that are entirely analogous to those described below with respect to row-wise correlations between stimuli. However, the value of *n* is typically large in reverse correlation experiments, and furthermore there is reason to believe that correlation between the rows has a more direct impact on the estimation quality, as outlined below.

**Figure 1.**
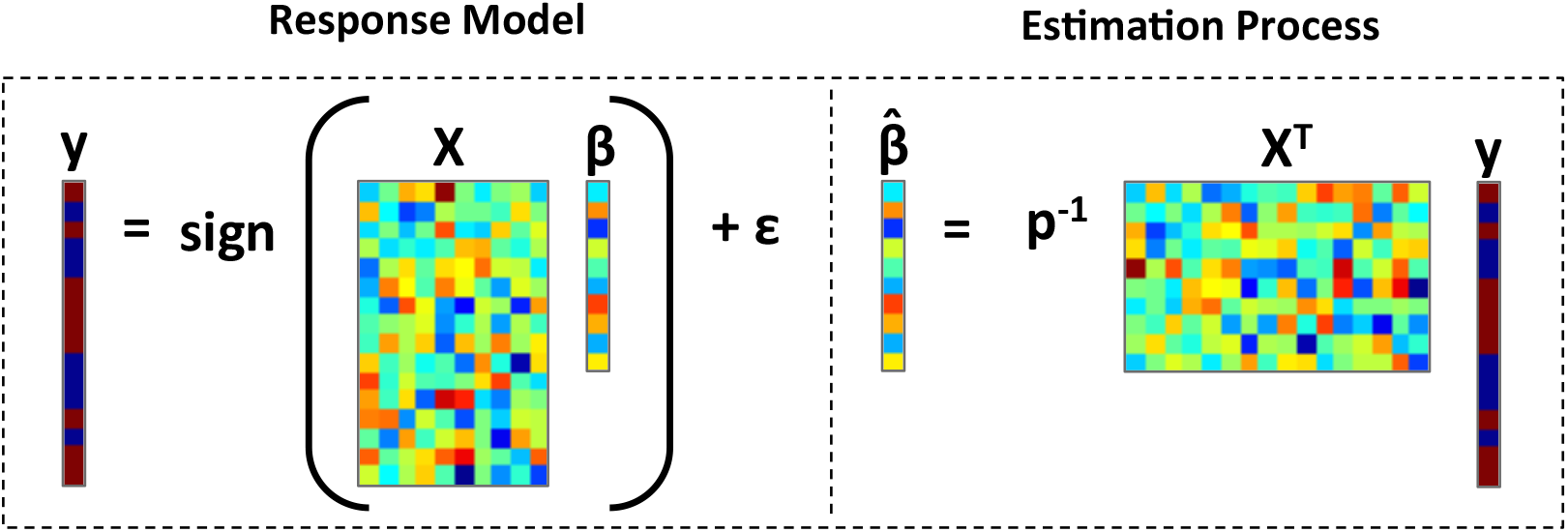
In reverse correlation, the vector of subject responses (y) is modeled as resulting from the multiplication of a latent representation vector (β) and a stimulus matrix (X). This can be thought of as calculating the similarity between the latent representation and a vector representation of each presented stimulus. To estimate the latent representation, the responses are regressed against the stimuli.

### Efficiency of Reverse Correlation

Crucial to improving the efficiency of reverse correlation is the ability to quantify estimation quality. Here, we quantify estimation quality by applying Pearson’ s product-moment correlation coefficient between the internal representation, *β*, and the estimate of that representation, 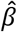, using the following equation:

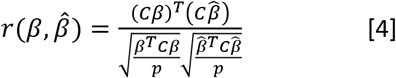

where 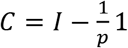 is the centering matrix and 1 is a matrix of all ones. Note that this metric of estimation quality assumes, as is typical in reverse correlation experiments, that internal representations encode only the relative values of the signal and not its overall magnitude. Given this metric of estimation quality, the goal of improving efficiency can be stated more precisely as maximizing the value of *r*(*β*, 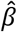) while limiting the number of trials *n*.

### Expected Correlation of Random Stimuli

In typical reverse correlation experiments, *n* stimuli of dimensionality *p* are randomly generated and presented to the subject in sequence. Here, we consider the case where all *n* stimuli are generated prior to initiation of the experiment as an *n*-by-*p* matrix *X*. If the stimuli are images, for instance, the rows of *X* represent images composed of *p* pixels, which can be reshaped into the desired two-dimensional format prior to presentation. Each element of *X* is drawn from a normal distribution with mean zero and variance one:

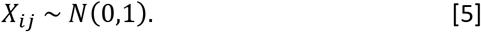

Equivalently, the stimuli are generated as the *n* rows of *X*, where each row is drawn from a *p*-variate normal distribution with mean zero and covariance matrix *V* = *I*:

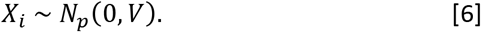

To examine the similarity of randomly-generated stimuli, we begin by considering the row-wise scatter matrix *S* of *X*, which contains the inner product between all pairs of stimuli. The matrix *S* is known to follow a Wishart distribution:

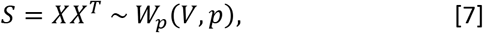

where *V* is a scale matrix and *p* is the degrees of freedom of the distribution. Accordingly, the mean of *S* is *pV* = *pI*, which has off-diagonal elements of zero, indicating that expected similarities between stimuli across experiments is nil. However, the variance of elements of *S* is

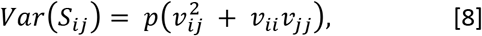

for elements *v_ij_* of *V*. Assuming once again that *V* = *I*, the above expression simplifies to

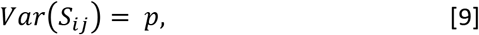

indicating that the similarities between stimuli will be expected to vary substantially between experiments, and therefore remain nontrivial for any given experiment.

Extending this analysis beyond the scatter matrix to the covariance matrix of *X* can help to clarify the expected magnitude of similarities between stimuli irrespective of their dimensionality. The covariance matrix can be written as:

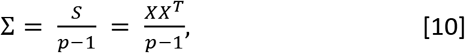

assuming for the sake of simplicity that the rows of X are mean-zero. One can find the variance of element of the covariance matrix, Σ*_ij_*, in the case where *V* = *I*, as:

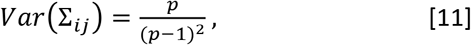

by observing that 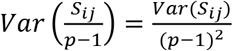. The variance of elements Σ*_ij_*, indicates, as above, that the similarities between stimuli are expected to be nontrivial for any given experiment, when those stimuli are composed simply of elements drawn from the Normal distribution. Examining the covariance matrix clarifies that this expectation is especially important when the dimensionality of the stimuli is low (i.e., *p* is small), which is often the case in reverse correlation experiments. The expected variance of elements *C_ij_*, and associated similarity between stimuli, can be eliminated through the process of whitening. Below, we describe how stimuli can be whitened, and provide a mathematical justification for why whitening is expected to improve estimation quality and the efficiency of reverse correlation.

## Method

### Whitening and Estimation Quality – Mathematical Justification

Our goal here is to show that whitening the rows of the n-by-p matrix of stimuli, *X*, will maximize the correlation between *β* and 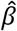. To clarify the role of *X*, we proceed to write Eq. 4 in terms of only *X* and *β*. Substituting values for 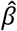 and *y* from the regression equations, stated above, the numerator can then be rewritten as 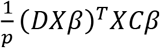. Using these same observations, the denominator can be rewritten as 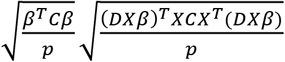. Together, these alterations yield the following equation:

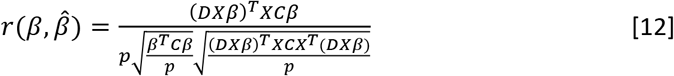

As mentioned above, we assume that internal representations encode only the relative values of the signal and not its overall magnitude. Therefore, it is reasonable to consider *mean*(*β*) = 0 and that ∥*β*∥_2_ = 1, which allows us to simply Eq. 12 further, becoming:

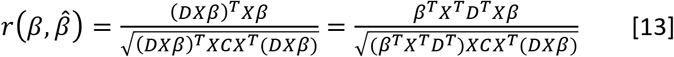

because *Cβ* = *β* and *β^T^Cβ* = 1 under the above assumptions, respectively. Again, the goal is to maximize the value of this equation for an arbitrary value of *β*, which can be done by maximizing the numerator and/or minimizing the denominator.

The numerator of Eq. 13 effectively compares the similarity, by way of taking the inner product, between *β* and each stimulus, and then sums the absolute values of those comparisons. This value is maximized when rows of X are equal to ±*β*, and therefore cannot be optimized without prior knowledge of the value of *β*. Assuming prior knowledge of the value of B is an approach often taken in the literature for improving the efficiency of reverse correlation experiments. However, such knowledge will always introduce estimation bias *a priori*.

The denominator of Eq. 13 clearly depends upon the similarity of rows of *X*, owing to the calculation of the centered row-wise scatter matrix *XCX*^T^, but is more difficult to analyze than the numerator. To simplify the analysis, we assume that for each row *i* of *X, mean*(*X_i_*) = 0 and that ∥*X_i_*∥_2_ = 1, both of which are similar to the assumptions made above in that they are consistent with the idea that internal representations encode only the relative values of the signal and not its overall magnitude. Given these assumptions, it can be easily verified that when *X* has been whitened with respect to its rows, meaning that *XCX^T^* = *I*, the denominator of Eq. 13 is equal to 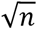. It can also be easily verified that when *X* is anti-white with respect to its rows – i.e., all off-diagonal elements of *XCX^T^* are equal to ±1 (e.g., *XCX^T^* = 1) – the value of the denominator in Eq. 13 is equal to 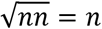. Therefore, using whitened stimuli is much more favorable than using stimuli that have the opposite statistical properties because 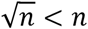.

The expected value of the denominator for typical stimuli – i.e., matrix X such that *X_ij_* ~ *N*(0, 1) – was estimated in this work through a series of numerical simulations. In each of these simulations, the value of the above denominator was evaluated for a randomly generated *X* and *β*~ *N*(0, 1). The values of *n* (the number of stimuli) and *p* (the dimensionality of the stimuli) were assigned to 8, 16, 32, 64, 128 or 256, such that all combinations of *n* and *p* were considered. For each unique combination of *n* and *p*, 1000 total simulations were conducted. It was found that, for a unique value of *n* and *p*, the mean value of the denominator was well-described by the formula 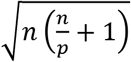. Note that the value of this formula exists between 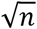 and *n* for most relevant values of n and p. It is higher than 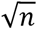 for all values of *n, p* ≥ 1, and lower than *n* for all values of *n* other than *n* ≫ *p*, or more specifically all values of 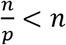, and then approximately equal to *n*. Furthermore, the denominator is equal to 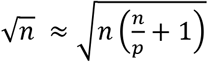 when *p* ≫ *n*, which is consistent with the expectation that rows with many elements (i.e., stimuli of high dimensionality) will be less covariant on average. Critically, the fact that the denominator value from this formula is higher than 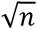 for all reasonable values of *n* and *p* confirms that whitening the matrix *X* can be expected to maximize *r*(*β*, 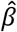), and therefore estimation quality.

### Whitening and Estimation Quality – Empirical Demonstration

To assess whether the theoretical efficiency improvements associated with whitening the stimuli could be observed empirically, a series of simulations were conducted in Matlab to assess estimation accuracy as a function of number of trials. The simulations were designed to follow Gosselin & Schyns (2003), in which one target of study was the internal representation of the printed letter ‘S’. In this study, three subjects completed 20,000 trials, in which subjects were shown random images (i.e., with pixel values drawn from a Bernoulli distribution) and asked to indicate, with a simple yes/no response, whether the image contained the letter ‘S’. Each subject’ s responses were used to generate an estimate using reverse correlation. One subject’ s estimated representation of ‘S’ (shown in Fig 3A) was used as the internal representation, *β*, in the simulations described here.

For each simulation, a normally-distributed random stimulus matrix of size *n*-by-*p* was generated as described in Eq. 5. This stimulus matrix was then either whitened (see Eq 15) or left unwhitened. Responses were generated using the assumed response generating process described in Eq. 1. Representation estimates were estimated using the typical regression-based reverse correlation procedure described in Eq. 3. Simulations were conducted over values of *n* ranging from 1000 to 5000 at increments of 1000. At each value of *n*, a total of 32 independent simulations were conducted, 16 with unwhitened stimuli, and another 16 with whitened stimuli.

For all simulations at a given value of *n*, the mean estimation accuracy and 95% confidence intervals were calculated separately for whitened and unwhitened stimulus matrices. As above, estimation accuracy was defined as *r*(*β*, 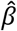).

### Whitening Procedure

For an unwhitened stimulus matrix *X_u_* of size n-by-p, the whitening matrix *W* may be defined as follows:

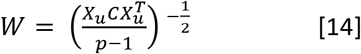

where *C* is the centering matrix. Other whitening matrices are possible (see, e.g., Kessy, 2015). This specific whitening procedure is sometimes called Mahalanobis whitening, or ZCA (zero-phase component analysis) whitening and can be seen as inverting the matrix square root of the row-wise covariance matrix of *X_u_*. Using the whitening matrix, one can calculate the matrix 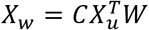, which is the data matrix *X_u_* with whitened rows.

Note that whitening in this way can become numerically difficult for very large values of n, due to difficulties inverting the relevant matrix. This can be overcome, and the range of possible values of *n* expanded, by introducing a slight bias to the diagonal of the matrix to be inverted of the form:

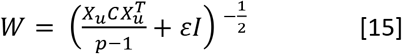

where the value of *ε* is small. This is conceptually similar to the procedure known as ridge regression. For the sake of consistency in comparing results, the procedure was implemented for all values of *n*, with *ε* = 0.001.

## Results

Figure 2 contains the simulation results, showing estimation accuracy, *r*(*β*, 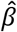) as a function of number of trials, *n* (mean and 95% CI). Accuracies using whitened and unwhitened stimuli are shown separately. Mean accuracy was found to be higher and variability in accuracy was found to be lower using whitened stimuli versus unwhitened stimuli at all values of *n*.

**Figure 2.**
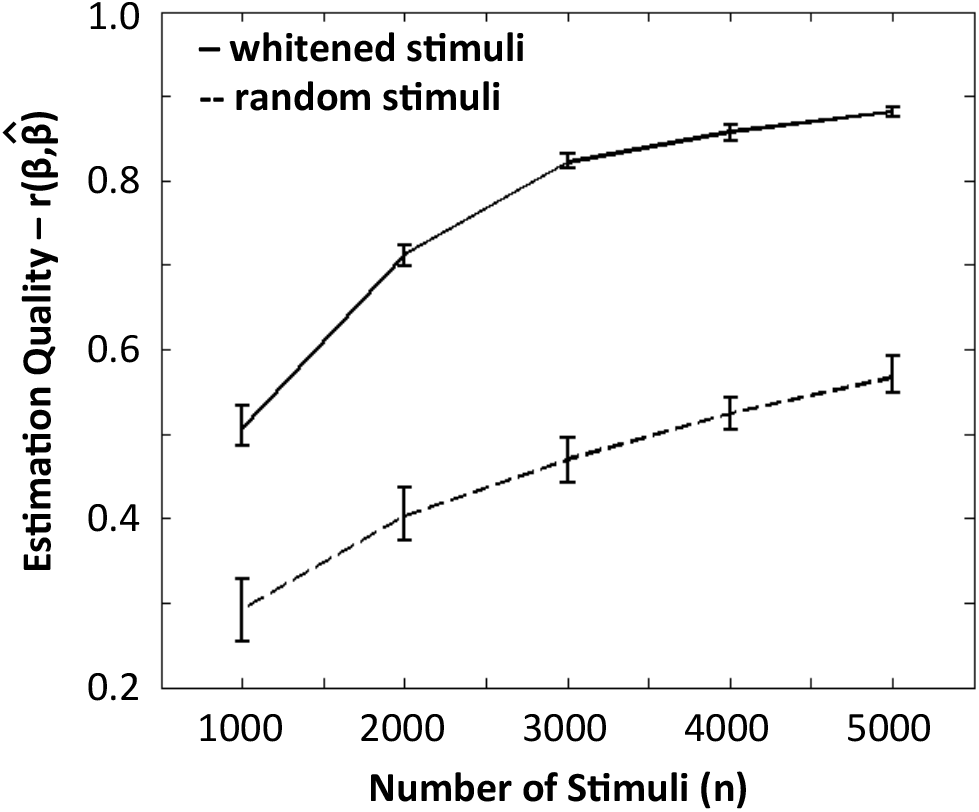
Estimation quality (mean and 95% CI) for both random stimuli (dashed line) and whitened stimuli (solid line) as a function of number of stimuli presented in a simulated reverse correlation experiment.

**Figure 3.**
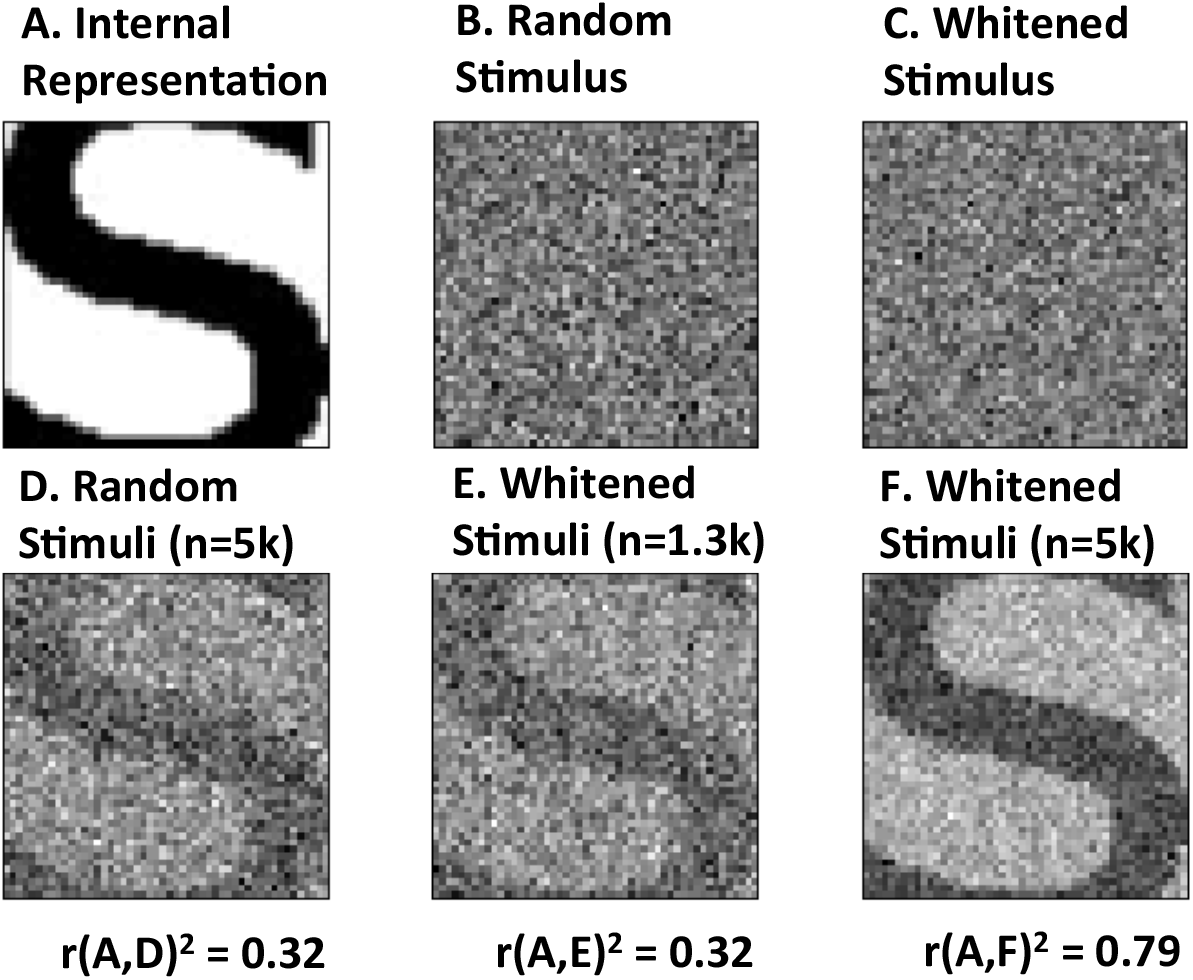
Comparison of reconstruction quality using conventional random stimuli and whitened stimuli. Example random/unwhitened stimuli and whitened stimuli are shown in B and C, respectively. Estimates of the template image A are shown in D-F, with the stimulus type and number of stimuli used (n) indicated above those images, and the correlation coefficient between the template and the estimate (r^2^, an indication of estimation quality) shown below.

Additional simulations (not shown) were conducted to determine the number of trials required, using unwhitened stimuli, to reach mean accuracy equivalent to 5k trials using whitened stimuli. The simulation procedure was repeated at increasingly higher values of *n*, in increments of 1k, until the estimation accuracy was equivalent to the accuracy obtained was at least *r* = 0.89. It was found that 40k trials were required to reach this level of accuracy, which represents an effective reduction in the number of trials of approximately 87.5%.

To facilitate a qualitative assessment of the estimation accuracy relative to both whitening and number of stimuli, example estimates using both unwhitened and whitened stimuli were randomly selected at two contrasting sample sizes (specifically, 1k stimuli and 5k stimuli). These examples were reshaped and plotted as images in Figure 3.

## Discussion & Conclusion

The mathematical justification provided above revealed that the two major sources of variability in estimation accuracy, as quantified by correlation between the estimate and the target representation, are (a) the degree of correlation among stimuli (the denominator in Eq. 13), and (b) the degree of correlation between the stimuli and the target (the numerator in Eq. 13). As such, lowering the degree of correlation among stimuli is expected in general to increase estimation accuracy. Whitening the stimuli is a process designed to accomplish exactly this goal, and should be expected, therefore, to improve estimation accuracy in reverse correlation.

Empirical results from the simulation study presented here indicate that the efficiency of reverse correlation is greatly improved by whitening stimuli before presenting them to subjects. For a given number of trials, estimation accuracy was observed to improve substantially when whitened stimuli were used, as compared to stimuli that were randomly generated and left unwhitened. This improvement was observed for all considered numbers of trials. Furthermore, when using whitened stimuli, the number of trials required to produce estimates of equivalent accuracy was substantially reduced, as well.

The empirical results also revealed that variance in estimation quality is sharply reduced by whitening stimuli. Again, the mathematical justification above revealed that the degree of random correlation among stimuli is a major source of variability in estimation accuracy. By eliminating any such correlation, whitening leaves only random correlation between the stimuli and the target as a source of variation in estimation quality.

Reverse correlation has the potential to uncover latent representations underlying perception and transform our understanding of perceptual mechanisms at various levels of investigation: neural, cognitive and psychological. However, in order for this potential to be fully realized, the fundamental inefficiency of reverse correlation paradigms must be overcome so that the breadth of its application may be increased. Whitening stimuli provides for more accurate estimates with fewer trials than simply using random stimuli, as in traditional approaches. Moreover, whitening does not impose any prior assumptions on the estimation process regarding the target representation. The dramatic improvements in efficiency demonstrated here can enable researchers to access the promise of reverse correlation by broadening its scope of application, allowing for studies to examine a wider array of representations within one individual, and also allowing deeper investigations into individual variability in, and potentially universal aspects of, perceptual representations.

